# Image-based deep learning reveals the responses of human motor neurons to stress and ALS

**DOI:** 10.1101/2021.04.08.439054

**Authors:** Colombine Verzat, Jasmine Harley, Rickie Patani, Raphaëlle Luisier

**Author notes:** Correspondence should be addressed to Rickie Patani and Raphaëlle Luisier. These authors contributed equally to this work (Rickie Patani, Raphaëlle Luisier). These authors contributed equally to this work (Colombine Verzat, Jasmine Harley).

## Abstract

Although morphological attributes of cells and their substructures are recognized readouts of physiological or pathophysiological states, these have been relatively understudied in amyotrophic lateral sclerosis (ALS) research. In this study we integrate multichannel fluorescence high-content microscopy data with deep-learning imaging methods to reveal - directly from unsegmented images - novel neurite-associated morphological perturbations associated with (ALS-causing) VCP-mutant human motor neurons (MNs). Surprisingly, we reveal that previously unrecognized disease-relevant information is withheld in broadly used and often considered ‘generic’ biological markers of nuclei (DAPI) and neurons (βIII-tubulin). Additionally, we identify changes within the information content of ALS-related RNA binding protein (RBP) immunofluorescence imaging that is captured in VCP-mutant MN cultures. Furthermore, by analyzing MN cultures exposed to different extrinsic stressors, we show that heat stress recapitulates key aspects of ALS. Our study therefore reveals disease-relevant information contained in a range of both generic and more specific fluorescent markers, and establishes the use of image-based deep learning methods for rapid, automated and unbiased testing of biological hypotheses.

## INTRODUCTION

Amyotrophic lateral sclerosis (ALS) is a relentlessly progressive and incurable neurodegenerative disease characterized by the loss of motor neurons (MNs). Key hallmarks of the disease include the mislocalization and accumulation of ubiquitously expressed RNA binding proteins (RBPs) from the nucleus to the cytoplasm including TAR DNA-binding protein of 43 kDa (TDP-43), Fused in Sarcoma (FUS) and Splicing factor Proline and Glutamine rich (SFPQ) proteins ^1–4^. While it is still unknown what drives pathological RBPs mislocalization and aggregation in ALS, alteration in liquid-liquid phase separation dynamics and functions have been proposed to underlie this process ^5–9^. RBPs are highly dynamic and have been shown to undergo changes in localization in response to various stressors ^10–16^. Notably mitochondrial and oxidative stress are additional recognized and robust phenotypes in ALS pathogenesis in vitro ^17^. The role of RBPs in ALS and cellular stress highlights that a diverse and complex interplay exists.

Cell shape and morphology are recognized readouts of a cell’s physiological state or phenotype ^18^. We previously reported common morphological descriptors that strongly discriminate ALS from control tissue at single cell resolution ^19^, further indicating that key information related to cellular state might be contained in cellular shape in ALS. Dystrophic neurites are common pathological features in ALS and disrupted synapse formation have been shown in valosin-containing protein (VCP) mutant human induced pluripotent stem cell (iPSC) cultures of MNs ^20^. Taken together, these studies suggest that the neuronal processes (collectively termed neurites or the ‘neuritome’) may be a good cellular subcompartment to reveal ALS pathomechanisms. However, neurites are challenging to study both in tissue sections (as the arborization of processes is largely lost during sectioning) and in vitro due to difficulty in accurate segmentation and association of neuronal processes with individual cells. Consequently, neuronal processes remain comparatively understudied in ALS and it is still unknown how and to what degree the neuritome is affected in ALS pathogenesis, whether ALS-related stress insults modify this compartment, or if cytoplasmic accumulation of RBPs in ALS MNs relates to other aberrant cellular phenotypes such as dystrophic neurites.

We previously generated a high-content imaging data-set of control and ALS-related VCP-mutant iPSC-derived MNs cultures co-labeled with a combination of three fluorescent markers, specifically: i) a nuclear-specific marker (DAPI), ii) a neuron-specific marker allowing to outline the neurites (βIII-tubulin), and iii) an antibody against one of five RBPs: TDP-43, SPFQ, FUS, heterogeneous nuclear ribonucleoprotein A1 (hnRNPA1) or heterogeneous nuclear ribonucleoprotein K (hnRNPK) ^16^. In our previous study we specifically analyzed the spatiotemporal responses of the aforementioned ALS-related RBPs to different stressors (oxidative, heat and osmotic). Here we propose to apply deep-learning methods to this rich imaging data-set to test in an automated fashion 1) whether aberrant cellular morphological phenotypes, including neuronal processes, associate with ALS pathogenesis; 2) whether these morphological phenotypes correlate to aberrant ALS-related RBP phenotypes, and 3) whether extrinsic stress insults in control MN cultures can recapitulate ALS phenotypic changes. Deep learning models such as Convolutional Neural Networks (CNNs) are now widely used to efficiently perform image classification and image segmentation ^21–25^. Such methods are able to analyze images without prior image segmentation, feature selection or human-directed training, and automatically extract features from raw data, removing significant bias from this process. Importantly CNN-based image classifier performance largely depends on whether sufficient information is contained in the provided set of images. DAPI and βIII-tubulin capture complementary and non-overlapping information related to the nuclear shape and neuronal morphology including the neurite respectively. We propose that comparing the performance of different classifiers trained with iterative combinations of fluorescent images can be used to identify which cellular compartment or specific RBPs is most affected between any two given culture conditions. Additionally we propose that the similitude in phenotypes between different MNs culture conditions can be quantified using the trained model predictions. We demonstrate the utility of this approach, which enables the discovery of novel phenotypes in ALS MN cultures and the identification of the relevant extrinsic stress condition that best approximates ALS pathogenesis. The advantage of our method is that it is highly versatile and can quickly guide the scientist towards the most promising hypothesis for further experimental validation. By providing our fluorescence microscopy raw images together with open-source implementations of the methods and trained models, we aim to allow other researchers to readily apply these methods and test additional hypotheses. In summary, we propose the use of deep learning methods to leverage the power of large image data-bases from ALS-related MN cultures to automatically and rapidly generate testable biological hypotheses, a method that could prove transformational in promoting innovative research directions, diagnostics and therapies.

## RESULTS

### Repurposing image-based deep learning methods to test biological hypotheses

We previously studied the spatiotemporal responses of ALS-related RBPs to different stressors in control versus ALS-related VCP-mutant iPSC-derived electrically immature MN cultures using image-based analysis (**Fig. 1A** and **Supplementary Table S1**) ^16^. These cultures were immunolabeled after one hour of exposure to oxidative stress, heat stress and osmotic stress, along with recovery timepoints from heat stress (two hours) and osmotic stress (one, two and six hours). Specifically a combination of three specific markers was used: a nuclear marker (DAPI), a neuronal marker allowing precise identification of neurites (βIII-tubulin), and an antibody against one of the following RBPs: TDP-43, SPFQ, FUS, hnRNPA1 or hnRNPK. Using this approach we generated a large-scale imaging data-set of 156,577 images (**Figs. 1B-D**). In our previous study we focused on nuclear-to-cytoplasmic ratio measurements of the aforementioned RBPs. Here we aimed to capitalise on the richness of information contained within this high-dimensional image data-set to test whether different MN stressors (including extrinsic stressors and endogenous ALS-causing mutations in the VCP gene) are characterised by detectable phenotypes in cellular compartments and/or RBP fluorescent images. Specifically we hypothesized that ALS-related phenotypic changes will be recapitulated by one of our aforementioned stress conditions. CNN-based classifiers are powerful deep learning models that can be trained to discriminate images from different conditions by identifying complex relationships between pixels. Here we trained the following 52 CNNs-based classifiers to recognize cellular phenotypes associated with i) ALS, ii) oxidative stress, iii) heat stress or iv) osmotic stress using 13 different combinations of immunolabeled images, ranging from DAPI fluorescent images only to the combination of three channels i.e DAPI, βIII and an RBP (**Fig. 1E** and **Supplementary Table S2**): 13 classifiers trained to discriminate untreated ALS from untreated control MN cultures (hereafter called ALS *classifiers*), 13 classifiers trained to discriminate untreated control MN cultures from control MN cultures exposed to oxidative stress (hereafter called OX *classifiers*), 13 classifiers trained to discriminate untreated control MN cultures from control MN cultures exposed to heat stress (hereafter called HS *classifiers*), and 13 classifiers trained to discriminate untreated control MN cultures from control MN cultures exposed to osmotic stress (hereafter called OSM *classifiers*). The CNN-based classifiers were obtained through transfer learning from MobileNetV2, which has been pre-trained using the ImageNet dataset^21^. The performance of each classifier was evaluated using the total area under the receiver operating characteristic (ROC) curve (AUC). AUC was calculated using 10-fold cross validation, training on 90% of the data-set, testing on the remaining 10% of the dataset, and repeating with 10 different train/test combinations (**Supplementary Table S3**). The 52 trained classifiers assigned class (e.g. ALS, stress) probabilities for all the ∼10 views from each cell culture (control versus ALS; untreated versus stressed; different time-points) that were then averaged to obtain a final per-culture classification probability (**Supplementary Tables S4-S7** and **Supplementary Fig. 1**).

**Figure 1.**
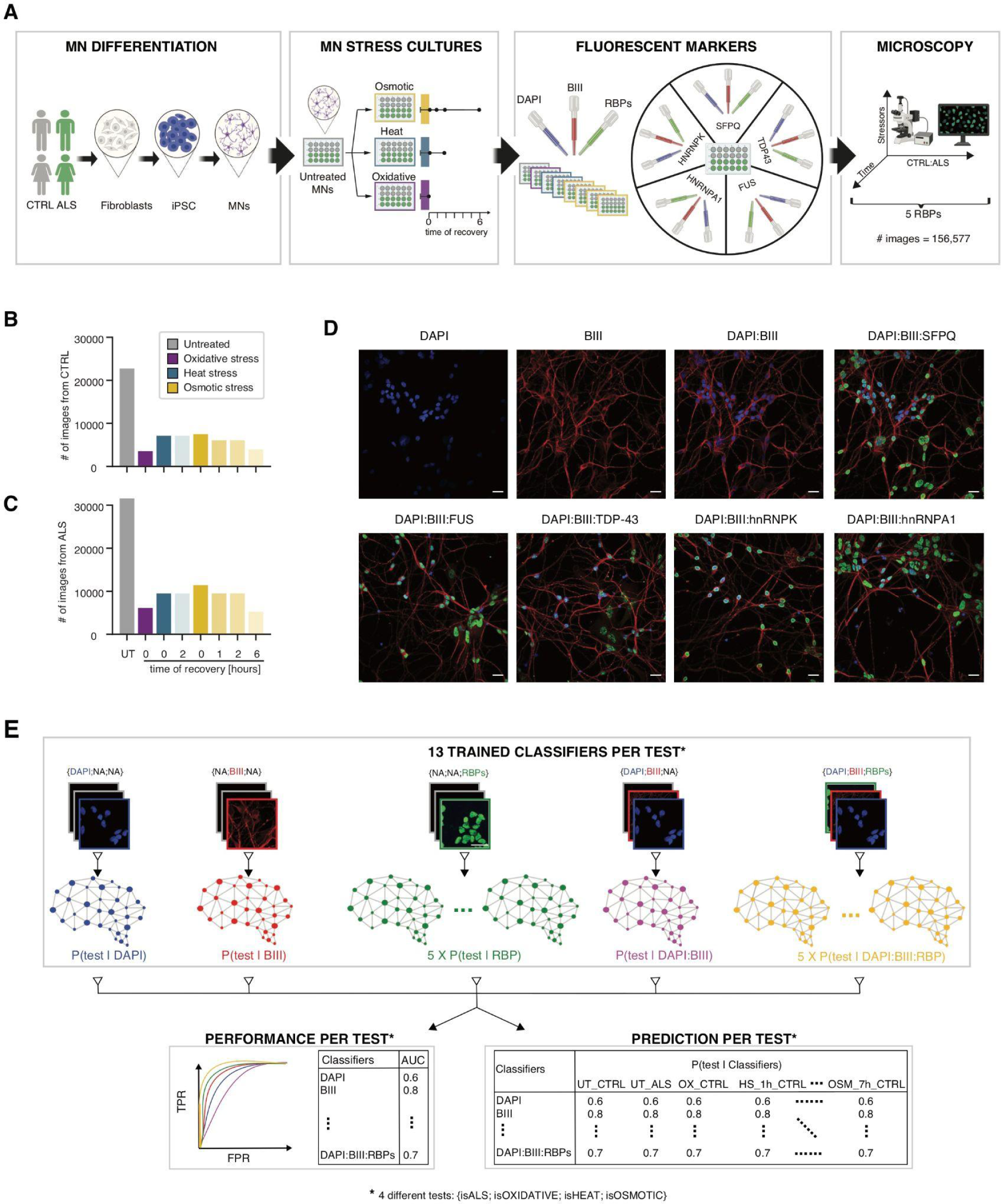
Overview of the high-dimensional immunofluorescent image data-set and paradigm to evaluate the relevance of markers in stress and ALS pathogenesis. (**A**) Experimental design for obtaining immunofluorescence microscopy images of motor neurons (MNs). Control (n = 3 cell lines) and VCP-mutant (n = 4 cell lines) induced pluripotent stem cells (iPSC)-derived MNs in different cellular stress (untreated, osmotic, heat, oxidative) and stress recovery (2h after heat stress, 1, 2 and 6h after osmotic stress) conditions were fluorescently labeled with DAPI, βIII-tubulin (BIII) and key ALS-linked RNA-binding proteins (RBPs) and then imaged, resulting in 156,577 images. (**B,C**) Total number of images in CTRL and ALS cell lines grouped by stress conditions: untreated (UT) in grey, oxidative (OX) in purple, heat (HS, 2h recovery) in blue and osmotic (OSM, 1h recovery, 2h recovery and 6h recovery) in yellow. (**D**) Representative images of nuclear marker DAPI, neuronal marker βIII-tubulin and ALS-linked RBPs in iPSC-derived motor neurons. Scale bars = 25μm. (**E**) 52 CNN-based classifiers have been trained in this study to discriminate 1) ALS from control MN cultures (isALS test), or untreated MN cultures from MN cultures exposed to 2) oxidative (isOXIDATIVE test), 3) heat (isHEAT test) and 4) osmotic stress (isOSMOTIC test) using 13 combinations of RGB images composed of different channels: either a single channel was used (DAPI, BIII or RBP), either two channels (DAPI:BIII) or three channels (DAPI:BIII:RBP) and pitch-black images were assigned to the unused channels (**Table S2**). For each of the four tests (siALS, isOXIDATIVE, isHEAT, isOSMOTIC), the 13 classifiers’ performance as obtained from the Area Under the Receiver Operating Characteristic Curve (AUC) were extracted and compared to uncover the importance of those markers in discriminating two conditions (**Tables S3**). Additionally 13 model predictions for each of the 4 tests have been extracted for each MN culture (**Tables S4-S7**).

Noting that the information content of images determines the performance of a CNN-based classifier to discriminate between conditions, we harnessed distinct fluorescent markers (DAPI, βIII-tubulin and RBPs) to capture different cryptic attributes that reveal cellular state. Against this background, we propose that the performance of the 13 different classifiers trained to identify a specific MN culture condition can reveal the relevant cellular compartment or RBP. During training, a classifier learns to identify a phenotype associated with a specific MN culture condition (ALS versus control; stressed versus untreated). Therefore, we propose that once trained, this classifier can be used to predict whether similar phenotypes are shared among different conditions. Consequently, we use image-based deep learning methods in two novel ways that are expected to greatly facilitate and accelerate the process of hypothesis testing in biology. In the following sections we first validate our approach by recapitulating previous findings. We next specifically demonstrate the utility of this approach by testing the following hypotheses: 1) ALS-causing VCP mutations result in previously unrecognized phenotypes contained within the information content of DAPI and/or βIII-tubulin fluorescence images alone; 2) addition of ALS-related RBPs immunofluorescence images will improve phenotype detection in an RBP-specific manner, and 3) conventional extrinsic stressors can recapitulate phenotypic aspects of ALS.

### Post-stress recovery of RBP- and neuritome-related phenotypes are closely correlated

Cell shape and morphology are recognized readouts of cell state or phenotype ^18^. Here we first sought to test whether oxidative, heat or osmotic stress are characterized by changes in cell shape. We analyzed and compared the performance of CNN-based models trained to discriminate images of untreated control MN cultures from images of stress-treated control MN cultures either using the DAPI staining only (hereafter named *stress|DAPI* classifier), either the β III-tubulin immunolabeling only (hereafter named *stress|BIII* classifier) or the combination of two fluorescent markers (hereafter named *stress|DAPI:BIII* classifier). As shown in **Fig. 2A**, *stress|DAPI* and *stress|BIII* classifiers outperform a random classifier (*AUC*_*DAPI*_ > 0. 5 and *AUC*_β*III*_ > 0. 5) across all stress conditions, indicating that DAPI and βIII-tubulin fluorescent images capture relevant information related to stressed MNs. This supports the hypothesis that MNs stressed with oxidative, osmotic or heat exhibit previously unrecognized phenotypic changes in both nuclear and neuritome compartments. Furthermore, comparing the performance of these classifiers revealed that the βIII-tubulin immunolabeling consistently leads to a significantly higher performance across all three stress conditions, suggesting that the compartment most affected by the all three stressors is the neuritome. Although we cannot rule out the possibility that the increase in model performance between *stress|DAPI* and *stress|BIII* is due to the larger surface occupied by the neuronal processes compared to the nuclei, the minor increase in model performances across the three stress conditions between *stress|BIII* and *stress|DAPI:BIII* supports the hypothesis that DAPI-stained images are not major contributors in these classifiers. The greater performance of *osm|DAPI* compared to *ox|DAPI* and *heat|DAPI* finally suggests larger nuclear-related changes upon osmotic stress compared to the other stress insults.

**Figure 2.**
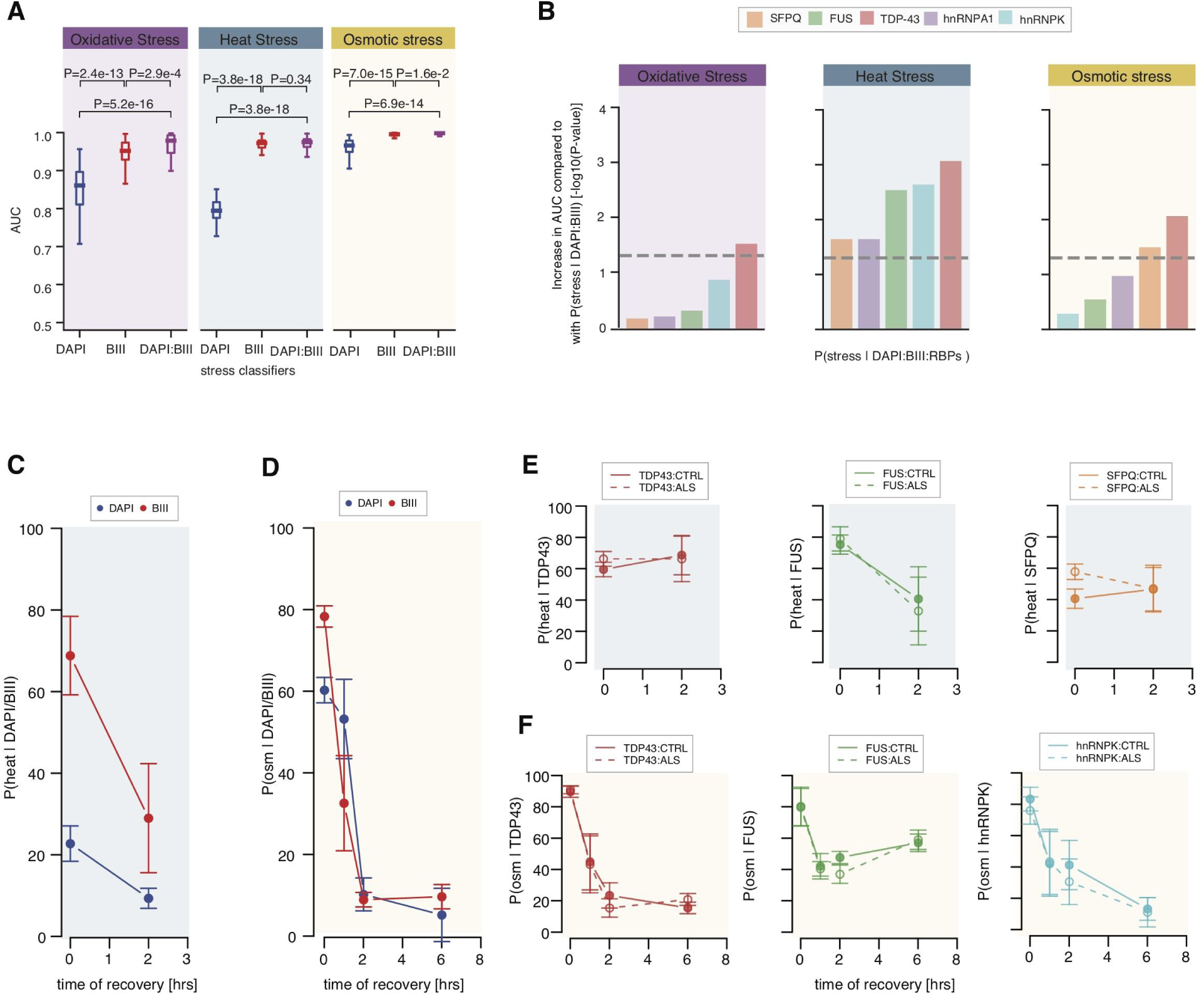
Different kinetics of stress recovery related to distinct cellular changes captured by CNN-based classifiers. (**A**) Boxplots showing the distributions of model performances as evaluated using the AUC for classifiers trained using DAPI, βIII tubulin (BIII) and the combination of DAPI and βIII tubulin markers, respectively, to discriminate untreated from stressed MN cultures. Each classifier was submitted to 10-fold cross-validation in 5 different subsets of the data, resulting in 50 points per classifier. Boxplots display the five number summary of median, lower and upper quartiles, minimum and maximum values. P-values obtained from a one-sided Mann-Whitney test. (**B**) Bar graphs representing the increase in performance as obtained from -log10(P-values) of one-sided Mann-Whitney test comparing the AUCs from the *stress|DAPI:BIII* classifier and the AUCs from individual *stress|DAPI:BIII:RBP* classifiers for oxidative, heat and osmotic stresses. (**C**) Effect size (mean ± standard errors) of *heat|DAPI*-based (blue) and *heat|BIII*-based (red) classifier predictions of control MN cultures after one hour exposure to heat stress and two hours of recovery from heat stress. Effect size of the treatment at each time-point is obtained using linear mixed effects analysis accounting for idiosyncratic variations due to cell lines and experiment bias. (**D**) Effect size (mean ± standard errors) of *osm|DAPI* (blue) and *osm|BIII* (red) classifier predictions of control MN cultures one hour after osmotic stress and one, two, and six hours after recovery from osmotic stress. (**E**) Same as (**C**) for *heat|TDP43*, *heat|FUS* and *heat|SFPQ* classifier predictions. Solid lines = control MN cultures. Dashed lines = VCP-mutant MN cultures. (**F**) Same as (**D**) for *osm|TDP43*, *osm|FUS* and *osm|SFPQ* classifier predictions. Solid lines = control MN cultures. Dashed lines = VCP-mutant MN cultures.

We previously showed that different stressors affect the localization of ALS-associated RBPs in control MNs ^16^. Thus we next aimed to test whether our image-based deep learning approach could shed light on the most relevant RBPs to each stress condition in order to replicate these previous findings. Examining the performances of the *stress|RBPs* models to discriminate untreated MN cultures from those exposed to osmotic-, heat- or oxidative-stress, and comparing these with the performance of *stress|DAPI* models revealed significantly higher performances of all five *stress|RBPs* models compared to *stress|DAPI* irrespective of the stress (**Supplementary Figs. 2A,B**). This result indicates that, although these RBPs mostly localize to the nucleus ^16^, their respective fluorescent images carry information beyond nuclear shape or texture as identified by DAPI. We next compared the performance of the *stress|DAPI:BIII:RBPs* models with the *stress|DAPI:BIII* models in each stress condition in order to test whether the integration of RBPs fluorescent images together with those of DAPI and βIII-tubulin enables the identification of additional stress-related phenotypes. This analysis revealed that TDP-43 significantly increases the ability of the classifier to identify MN cultures under oxidative stress (**Fig. 2B**). While CNN-based models are not suited to specifically address the subcellular localization of RBPs, our finding that TDP-43 images *in conjunction with nuclear and neurite fluorescent markers* enable relevant oxidative-stress related phenotypic information to be captured, suggests that TDP-43 exhibits changes in localization upon oxidative stress. This result is consistent with our prior finding that TDP-43 - but not the other four RBPs analyzed - exhibits a reduction in nuclear-to-cytoplasmic ratio upon oxidative stress ^16^. We also find that all five *stress|DAPI:BIII:RBPs* perform significantly better than *stress|DAPI:BIII* to discriminate untreated from heated MN cultures. Furthermore, we find that the most informative RBPs to heat stress are TDP-43, FUS and hnnRNPK. While in our previous study we detected significant reduction in nuclear-to-cytoplasmic ratio for TDP-43 and FUS upon heat stress, we can speculate that the present approach captures more subtle changes beyond the previously studied cellular relocalization that could explain the detected relevance of hnRNPK to heat stress. Finally, while we previously found that all five RBPs exhibit nuclear-to-cytoplasmic relocalization upon osmotic stress, here we find that TDP-43 and SFPQ immunolabeling only contribute to significantly increase the *stress|DAPI:BIII* performance to identify MN cultures under osmotic stress. It is however important to note that *osm|DAPI:BIII* exhibits an AUC of ∼1.0, implying that a significant improvement is difficult to achieve in this case and that in the case of osmotic stress, this analysis may underestimate the contribution of the RBP immunolabeling. Altogether these results indicate that the performance of a classifier is a reliable approach to prioritize which RBPs are most relevant to a specific cell culture condition.

Next the extent of recovered cellular compartment- and RBP-related phenotypes after heat and osmotic stress were assessed using linear mixed effects analyses of the individual classifier predictions, accounting for idiosyncratic variations due to either individual cell lines or experiments. As shown in **Fig. 2C**, two hours after recovery from heat stress, the nuclear compartment has fully recovered, as predicted by *heat*|DAPI, while the neuritome compartment still exhibits some degree of aberrant phenotype, as predicted by *heat*|BIII. As opposed to heat stress, the nuclear compartment takes longer to recover after osmotic stress compared to the neuritome compartment, however both compartments exhibit full recovery 6 hours after treatment (**Fig. 2D**). Next looking at the RBP-related phenotypes, we find large heterogeneity in their predicted recovery pattern after both heat and osmotic stresses, with no complete recovery for any of the analysed RBPs two hours after heat stress (**Supplementary Figs. 2C,D**) and long-term effects for several RBPs after osmotic stress (**Supplementary Figs. 2E,F**). In particular we find that two hours after heat stress, MN cultures still exhibit high *heat|TDP-43* and *heat|SFPQ* model predictions and lower (albeit still elevated) *heat|FUS* model prediction (**Fig. 2E**). The results indicate that the TDP-43- and SFPQ-related phenotypes are still present at this stage, and that the FUS-related phenotype is only partially resolved, partly reflecting on our previous study, where we did not detect reconstitution of nuclear TDP-43 and FUS to basal levels following 2 hours of recovery from heat stress ^16^. Our previous study also revealed slower nuclear relocalization dynamics for TDP-43 and FUS after osmotic stress, with FUS exhibiting exceptionally aberrant nuclear-to-cytoplasmic distribution as long as 6 hours post-stress ^16^. Here we find that TDP-43-related phenotype is fully resolved 2 hours after treatment while FUS-related phenotype is not resolved 6 hours after treatment (**Fig. 2F**). We also find delayed hnRNPK-related phenotype recovery. Notably we find that the recovery kinetics for most RBPs after both heat and osmotic stress correlate over time with the neuritome-related phenotype, suggesting that changes in neuritome relate to change in RBP-related phenotype or vice-versa. Finally, and in line with our previous study, we do not find any major difference between control and ALS-related VCP mutant MNs cultures in their response to stress (**Supplementary Figs. 2D,F**). While these results at least in part recapitulate our previous findings, thereby confirming the validity of our approach, it is important to note that the trained classifiers do not necessarily capture a phenotype related to the previously studied nuclear-to-cytoplasmic relocalization and that this may relate to more complex cellular response. Altogether these results indicate that the performance of a classifier is a reliable approach to prioritize which RBPs are most relevant to a specific cell culture condition and that the CNN-based method can, at least in part, reproduce previous results showing slower TDP-43 and FUS relocalization dynamics following heat and osmotic stress ^16^ that in some cases these might relate to changes in neuritome.

### Heat stress-related changes in the MN neuritome resemble those occurring in ALS

We previously reported common morphological descriptors that strongly discriminate ALS from control control tissue at the single cell level ^19^, indicating that key information related to ALS cellular state might be contained in cellular shape. Having found that our approach is suitable to reproduce prior findings related to stress in MNs, we next sought to test whether ALS-related VCP-mutant MNs are characterised by changes in cellular shape in the nucleus, neuritome or the combination of both. This was achieved by comparing the performances of CNN-based classifiers trained to discriminate images of untreated control MN cultures from images of VCP-mutant MN cultures either using the DAPI staining only (hereafter named *ALS|DAPI* classifier), either the βIII-tubulin immunolabeling only (hereafter named *ALS|BIII* classifier) or the combination of two markers (hereafter named *ALS|DAPI:BIII* classifier). As shown in **Fig. 3A**, both ALS:DAPI and *ALS|BIII* classifiers outperform a random classifier (*AUC_DAPI_* > 0. 5 and *AUC_βIII_* > 0. 5), indicating that, similarly to stress-related conditions, both compartments exhibit phenotypic changes associated with VCP mutation. Further comparing the performances of these classifiers revealed that the inclusion of the βIII-tubulin immunolabeling leads to consistently significantly higher performance (*AUC_DAPI+βIII_* = 0. 85> *AUC_βIII_* = 0. 83> *AUC_DAPI_* = 0. 65), indicating that the compartment most affected by VCP mutation at this early disease stage is the neuritome. Notably the consistently high model predictions among the four mutant cells lines according to both classifiers confirms the absence of experimental or cell line bias, as further confirmed by linear mixed model analysis (**Fig. 3B**). Additionally, the significant correlation between the *ALS|DAPI* and *ALS|BIII* model predictions for each MN culture further indicates that the ALS-related changes identified by these classifiers in the nucleus or the neuritome respectively co-occur in the same MNs cultures (Pearson correlation coefficient = 0.67 and *P*=2.93e-33; **Fig. 3C**).

**Figure 3.**
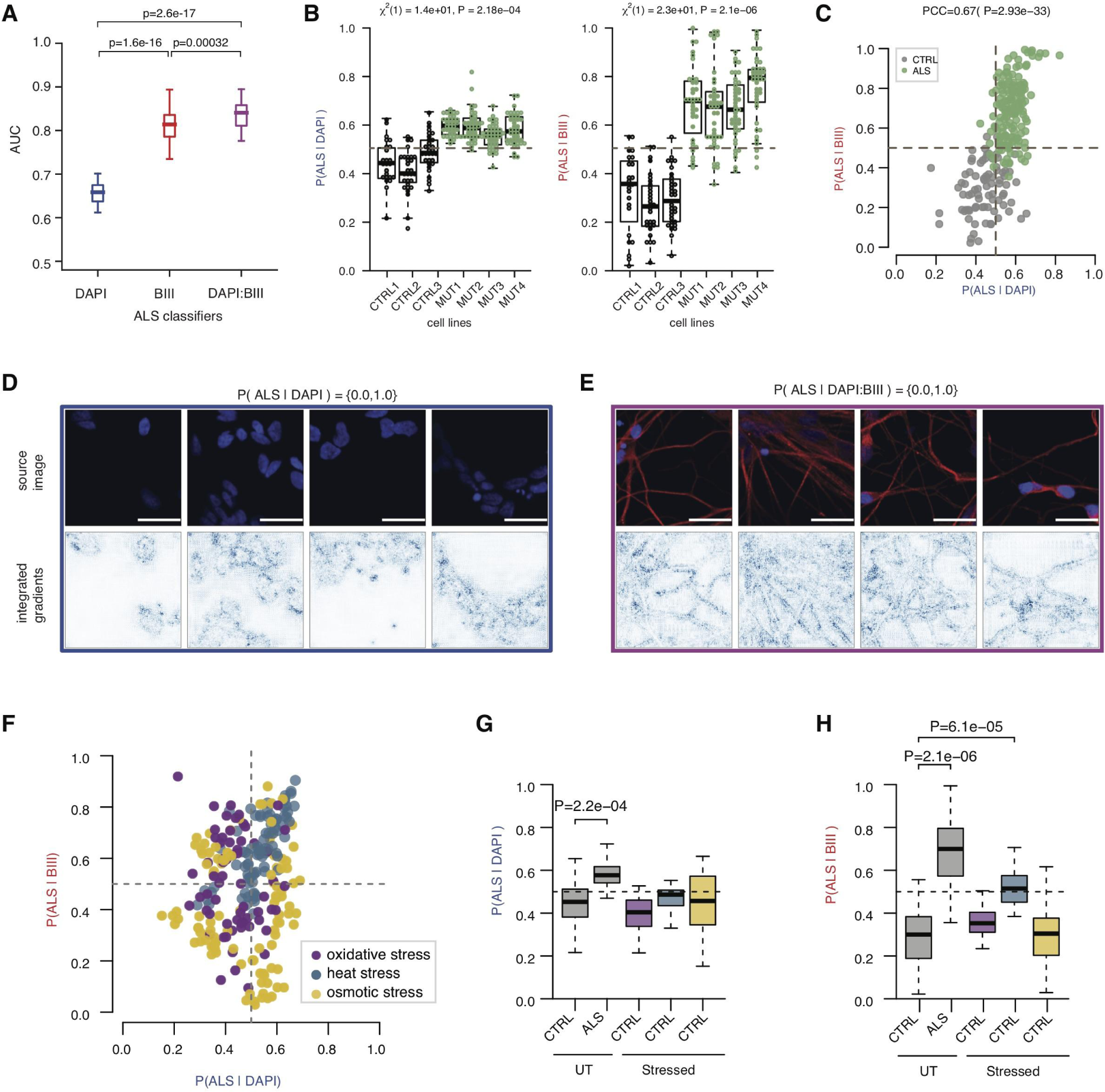
Heat stress-related changes in the MN neuritome resemble those occurring in ALS. (**A**) Boxplots showing the distributions of model performances as evaluated using the AUC for classifiers trained using DAPI, βIII-tubulin (BIII) and the combination of DAPI and βIII-tubulin markers to discriminate control from VCP-mutant MN cultures. Each classifier was submitted to 10-fold cross-validation in 5 different subsets of the data, resulting in 50 points per classifier. P-values are from a one-sided Mann-Whitney test. Data shown as in Fig. 2A. (**B**) Distributions of the *ALS|DAPI* (*left*) and *ALS|BIII* (*right*) model predictions for the individual MN cultures originating from 3 control cell lines (*grey dots*) and 4 VCP-mutant cell lines (*green dots*). Linear mixed effects analysis of the relationship between each model prediction and VCP mutation to account for idiosyncratic variation due to cell line or experiment differences. VCP mutation significantly increases *ALS|DAPI* predictions [χ^2^(1) = 14 and *P* = 2. 28*e*^-04^] by about 0. 13 ± 0. 023 (standard errors), and *ALS|BIII* predictions [χ^2^(1) = 23 and *P* = 2. 1*e*^-06^] by about 0. 4 ± 0. 033 (standard errors). (**C**) Scatter plot of *ALS|DAPI* and *ALS|BIII* model predictions on individual control and VCP-mutant MN cultures. Grey = control MN cultures. Green = VCP-mutant MN cultures. PCC = Pearson Correlation Coefficient. (**D,E**) Randomly selected images with high model prediction according to the *ALS|DAPI* (D) and *ALS|DAPI:BIII* (E) classifiers. Images shown are the original image (*upper*) and the corresponding attribution magnitude image overlayed on the original image (*lower*). The magnitudes range from 0 (white), indicating no contribution of the pixel, to 1 (blue), indicating the strongest contribution of the pixel to the model prediction. Scale bars = 25μm. (**F**) Scatter plot of the *ALS|DAPI* and *ALS|BIII* model predictions on individual control MN cultures one hour after oxidative, heat and osmotic stress. Magenta = MN cultures one hour after oxidative stress. Blue = MN cultures one hour after heat stress. Yellow = MN cultures one hour after osmotic stress. (**G**) Boxplots showing the distributions *ALS|*DAPI model predictions on untreated control and VCP-mutant MN cultures, and control MN cultures one hour after oxidative, heat and osmotic stress. Magenta = MN cultures one hour after oxidative stress. Blue = MN cultures one hour after heat stress. Yellow = MN cultures one hour after osmotic stress. Stress treatment effect analysis on model prediction obtained using linear mixed effects analysis. P-value is indicated when significant. (**H**). Same as (G) for *ALS|*BIII model.

We next aimed to understand what information is used by these ALS classifiers to discriminate images from control and VCP-mutant MN cultures. Integrated gradients (IG) is one popular approach for CNN model interpretation enabling the visualisation of the relevant pixels for a specific image that contribute to its classification ^26^. Looking at the IGs of randomly selected images with high *ALS|DAPI* model predictions showed relevant pixels mostly overlapping with the outline of the nuclei, with some contribution from the pixels located inside the nuclei (**Fig. 3D**). This indicates that the ALS-related phenotype identified by the *ALS|DAPI* classifier primarily relates to the nuclear shape (including the size) rather than to other DAPI-related measurements such as texture or intensity. Next looking at the IGs of randomly selected images with high *ALS|DAPI:BIII* model predictions showed relevant pixels primarily located at the edges of the neurites, indicating that relevant information mostly arises from the outline of the neurites rather than from the texture or the intensity of the βIII-tubulin immunolabeling (**Fig. 3E**). Altogether these results indicate the network of neurites carries most ALS-related phenotype information.

Mitochondrial and oxidative stress are recognized and robust phenotypes in ALS development in vitro, and thus *in vitro* models of cellular stress are important tools to investigate ALS disease^17^. However it remains unknown which type of cellular stress is most physiologically relevant to study ALS pathogenesis. Thus we next sought to test whether heat, osmotic or oxidative extrinsic stress insults induce similar nuclear and/or neuritome-related phenotypic changes in control MNs cultures as those captured by *ALS|DAPI* and *ALS|BIII* classifiers. First looking at the scatter plot of these two model predictions for individual MN cultures color-coded according to the stress conditions showed that heated MN cultures consistently score high according to both classifiers forming a coherent cluster, as opposed to oxidative and osmotic conditions (**Fig. 3F**). We next quantified the extent of ALS-related nuclear versus neuritome phenotypes induced by each individual treatment (oxidative, heat, osmotic) by analysing the *ALS|DAPI* or *ALS|BIII* model predictions of individual stressed MN cultures using linear mixed modelling (see **Materials and Methods**). This analysis revealed that heat stress, and to some extent osmotic stress, induces a minor however non-significant ALS-related nuclear phenotypic change in control MN cultures, as indicated by the increase in *ALS|DAPI* model predictions for these two MNs cultures (**Fig. 3G**). This is in contrast with the neuritome compartment which *ALS|BIII* model predicts significant phenotypic changes in heated MNs cultures only (**Fig. 3H**). Altogether these results indicate that the neuritome is the compartment most affected by ALS-related VCP mutation and that heat stress induces similar neurite-associated changes in control MN cultures.

### FUS immunolabeling best captures ALS-related phenotypes that are recapitulated by heat stress

Two-thirds of RBPs are expressed in a cell type-specific and temporally regulated manner ^27^. While previous studies showed RBP-specific mislocalization in various ALS models^2–4, 28, 29^, it remains unknown which RBP is most predictive of ALS at a particular disease stage. Thus we next sought to test whether different ALS-related RBPs capture distinct ALS-related information in VCP-mutant MN cultures by analyzing the model performances of ALS classifiers trained with images immunolabeled with antibodies against TDP-43, FUS, SFPQ, hnRNPK or hnRNPA1 only (hereafter named *ALS|RBPs* classifiers). Since all five aforementioned RBPs exhibit predominant nuclear localization ^16^, we first tested whether *ALS|RBPs* classifiers exhibit significant improvement compared to the *ALS|DAPI* classifiers. This analysis showed that all 5 *ALS|RBP* classifiers outperforms *ALS|DAPI* classifier (*AUC_ALS|RBPs_* > *AUC_ALS|DAPI_*), ruling out the possibility that the phenotypic changes captured by these classifiers simply overlap with those identified by the *ALS|DAPI* classifier and indicating that they identify ALS-related phenotypes beyond changes in the nuclear shape (**Fig. 4A**). Comparing their individual performances further revealed large differences in the individual RBP-based classifiers’ ability to discriminate ALS from control MN cultures, with *ALS|TDP-43* exhibiting the best performance and *ALS|SFPQ* the least (*AUC_ALS|SFPQ_* = 0.7 < *AUC_ALS|hnRNPK_*= 0. 73< *AUC_ALS|FUS_* = 0. 79 < *AUC_ALS|hnRNPA1_*= 0. 85 <*AUC_ALS|TDP43_* = 0. 9). Examining the IGs for randomly selected images with high *ALS|RBP* model predictions indicated that the relevant pixels in all five *ALS|RBPs* classifiers are excluded from the nuclear areas as opposed to the most relevant pixels of the *ALS|DAPI* classifier that are most commonly localized at the inner nuclear membrane or inside the nucleus (**Fig. 4B**). This demonstrates that the better the performance of the classifier, the less relevant the intranuclear pixels. For example relevant pixels in the *ALS|TDP-43* classifier are fully excluded from the nuclear area. Altogether these results suggest that the different performance of *ALS|RBPs* classifiers in identifying ALS MNs cultures result from distinct RBPs localization rather than nuclear shape. We previously showed that considering DAPI and βIII-tubulin together significantly increases the performance of both ALS|DAPI and ALS|BIII classifiers. We next analyzed how the performance of the ALS classifier would change by adding a third channel composed of RBPs immunolabeling and compared the performances of the *ALS|DAPI:BIII* classifier with the 5 *ALS|DAPI:BIII:RBP* classifiers. This analysis indicates that all five RBPs significantly increase the ability of the trained CNN to discriminate VCP-mutant from control MN cultures, however while hnRNPK leads to the most modest improvement, TDP-43 and FUS immunolabeling lead to the highest increases in classification performance (**Supplementary Fig. 4A**). These results support the hypothesis that TDP-43 and FUS exhibit changes in localization that are detected by these classifiers. Indeed, our prior studies have shown that in the same experimental model used here and at the same development stage, TDP-43 and FUS mislocalization already occur ^16, 20, 30^, thus we can expect similar RBP mislocalization. These results further suggest that all five RBPs exhibit mislocalization at different degrees which previous studies could not demonstrate possibly due to lower sensitivity.

**Figure 4.**
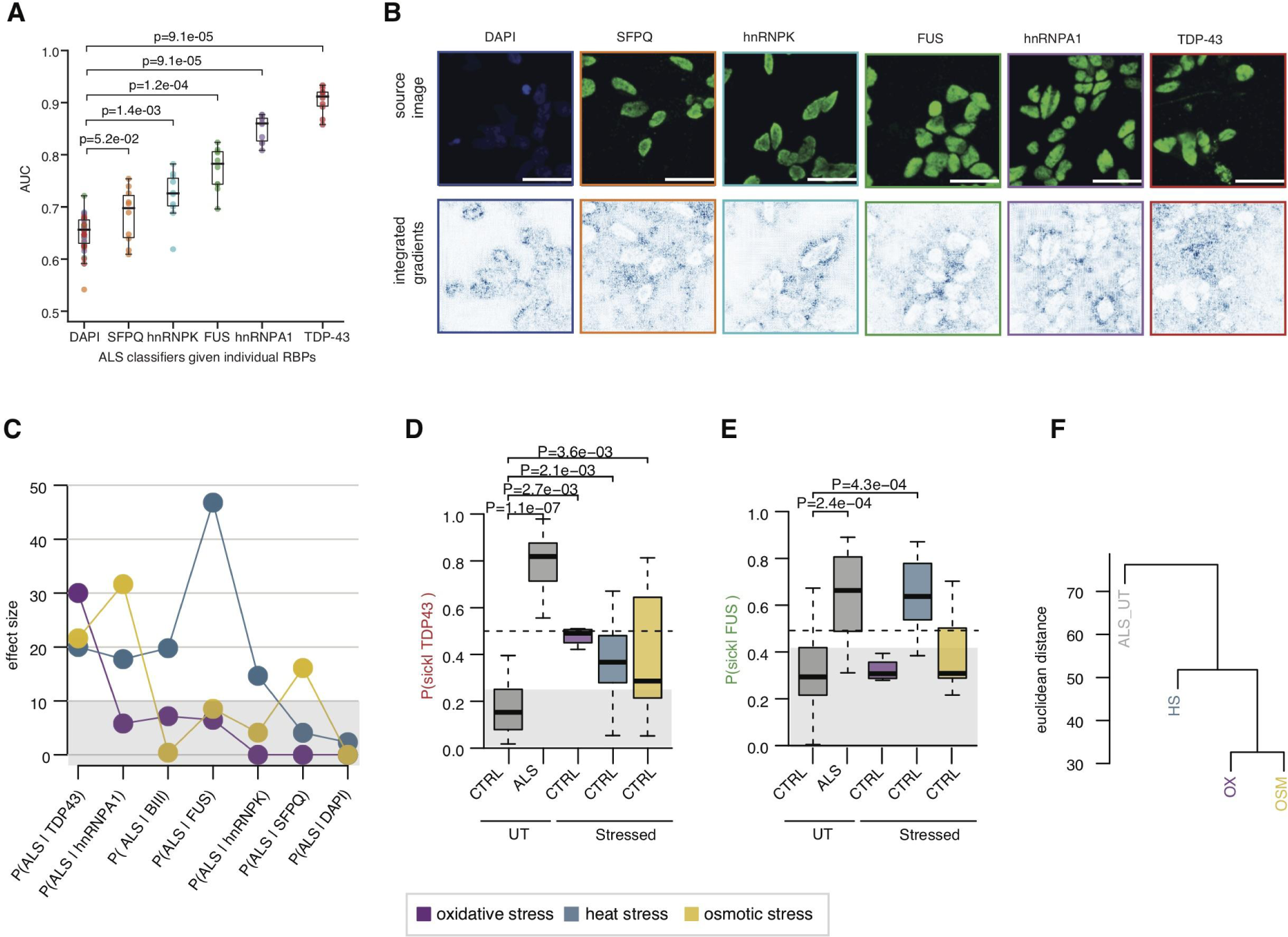
Heat stress as the most physiologically relevant stress condition to study ALS pathogenesis. (**A**) Boxplots showing the distributions of model performances as evaluated using the AUC for classifiers trained using the ALS-related RBPs markers SFPQ, hnRNPK, FUS, hnRNPA1 and TDP-43 to discriminate control from VCP-mutant MN cultures, and compared to *ALS|DAPI* classifier performance. Each classifier was submitted to 10-fold cross-validation in 5 different subsets of the data, resulting in 50 points per classifier. P-values are from a one-sided Mann-Whitney test. Data shown as in Fig. 2A. (**B**) Randomly selected images with high model prediction according to the *ALS|DAPI* and *ALS|RBPs* classifiers. Images shown are the original image (*upper*) and the corresponding attribution magnitude image overlayed on the original image (*lower*). The magnitudes range from 0 (white), indicating no contribution of the pixel, to 1 (blue), indicating the strongest contribution of the pixel to the model prediction. Scale bars = 25μm. (**C**) Comparisons of the effect sizes of each stress treatment on control MNs cultures *ALS|DAPI,* ALS|BIII and the 5 *ALS|RBPs* model predictions one hour after treatment. Magenta = oxidative stress. Blue = heat stress. Yellow = osmotic stress. Stress treatment effect analysis on model prediction obtained using linear mixed effects analysis. (**D**) Boxplots showing the distributions *ALS|*TDP-43 model predictions on untreated control and ALS MN cultures, and control MN cultures one hour after oxidative, heat and osmotic stress. Magenta = oxidative stress. Blue = heat stress. Yellow = osmotic stress. Stress treatment effect analysis on model prediction obtained using linear mixed effects analysis. P-values obtained from linear mixed models are indicated when significant. (**E**) Same as (D) however for *ALS|*FUS. (**F**) Unsupervised hierarchical clustering of treatment effect sizes on *ALS|DAPI,* ALS|BIII and the 5 *ALS|RBPs* model prediction groups. Oxidative and osmotic stresses cluster together while heat stress is the closest to the ALS group. Euclidean distance and ward clustering.

We next sought to test whether the ALS-related changes identified by the individual *ALS|RBPs* models are recapitulated in any of the extrinsic stressor cultures conditions. We first considered the effect size, as obtained by linear fixed effect analysis (**Materials and Methods**), of each extrinsic stressor culture condition on the predictions of the seven ALS classifiers (ALS|DAPI, ALS|BIII, *ALS|hnRNPK, ALS|SFPQ, ALS|FUS, ALS|hnRNPA1, ALS|TDP-43*) trained using a single channel. This revealed that oxidative and osmotic stress recapitulate only one or two ALS-related phenotypes out of the seven captured by the ALS classifiers. However, heat stress induced more than a 10% increase in ALS prediction across 5 out of 7 classifiers (**Fig. 4C**). *ALS|TDP-43* classifier, which performs best in ALS MN classification, is indeed the unique model which leads to significant, however modest, model prediction across the three stress conditions (**Fig. 4D** and **Supplementary Figs. 4B,C**). Additionally we find that heat stress is the unique condition which leads to similarly high disease model prediction in control MN cultures and in untreated VCP-mutant MN cultures, given the *ALS|FUS* and *ALS|hnRNPK* classifiers (**Fig. 4E** and **Supplementary Figs. 4B,C**). Hierarchical clustering of the untreated ALS MN cultures together with the three stress conditions according to the effect size of each classifier (euclidean distance and Ward clustering) eventually confirmed that heat stress induces overall the most similar cellular changes to ALS (**Fig. 4F**). Altogether these results confirm that MNs exposed to heat stress most closely resemble ALS cells with respect to phenotypes captured by the majority of ALS classifiers. Additionally, it shows that while TDP-43 is the RBPs that carries the strongest information related to ALS, it is the FUS immunolabeling which captures most similar phenotypes between heat stress and ALS MNs cultures.

## DISCUSSION

In this study, we combine multichannel fluorescence high-content microscopy data with deep-learning imaging methods to unveil - directly from unsegmented images - novel neurite-associated morphological perturbations. This approach can be used to leverage existing high content imaging datasets to gain new phenotypic insight into the original biological questions asked, as established by this study. We uncovered a surprising degree of previously unrecognized disease-relevant information in broadly used and often considered ‘generic’ biological markers of nuclei (DAPI) and neurons (βIII-tubulin) in the context of our human stem cell model of VCP-related ALS. Additionally, we reveal changes associated with ALS-related RBP immunofluorescence imaging that can be captured in VCP-mutant MN cultures. We were also able to systematically examine whether heat, oxidative or osmotic stress induce similar modifications that could therefore reinforce their utility in modeling aspects of MN dysfunction in ALS. Our study establishes the use of CNN-based methods for rapid, automated and unbiased testing of biological hypotheses.

CNN-based methods are now widely used for image classification and segmentation, and have been successfully applied to medical imaging data for disease detection and prediction ^31–33^. Here CNN-based image classifiers have been trained to identify stress- and ALS-related phenotypic changes in unsegmented images of multiple MN cultures. We further showed that the performance of such classifiers is a reliable approach to prioritize which RBPs are most relevant to a specific cell culture condition, although refined analysis will be required to interpret their precise relevance to the underlying disease/stress process. CNN-based classifiers face challenges with interpretability and are not suited to specifically address the subcellular localisation of RBPs as previously described with conventional methods based on image segmentation ^16, 30^. Nevertheless here we showed that the phenotypes identified in the DAPI or βIII-tubulin fluorescent images are indeed contained in the outlines of the nuclei and in the edges of the neurites respectively. Furthermore, we could demonstrate that training a classifier with fluorescent images of a given RBP *in conjunction with nuclear and neurite fluorescent markers* enables the recapturing of previously found phenotype related to RBP cellular localization. Specifically we could reproduce previous results showing TDP-43 and FUS mislocalization in ALS iPSC-derived MNs ^3, 20, 30^. Additionally our study suggests that SFPQ, hnRNPA1 and hnRNPK also exhibit mislocalization however at different degrees. Notably, hnRNPA1 is a component of RNA transport granules in neurons ^34^ and we can speculate that the extent of cytoplasmic relocalisation for this primarily nuclear RBP ^16^ may be too subtle to be captured by analysing its nuclear-to-cytoplasmic ratio.

DAPI and βIII-tubulin are often considered ‘generic’ biological markers and the usage of their fluorescent images are most often intended for nuclear or neuronal segmentation. Our study uncovers previously unrecognized disease-relevant information that is contained within DAPI and βIII-tubulin fluorescent images. Given that DAPI and βIII-tubulin are broadly used markers, and given that our method can be easily implemented in other biological conditions, future studies might use DAPI and βIII-tubulin fluorescent images from different biological systems and experimental paradigms to reveal innovative research directions. For example, this approach could be useful in interrogating the presence and onset of aberrant cellular morphologies in time course experiments and across neurological disorders.

Morphological attributes of cells and their substructures are recognized readouts of physiological or pathophysiological states ^18^, however these have been relatively understudied in ALS research. Here we demonstrate that the neuritome compartment exhibits aberrant phenotypes in ALS pathogenesis, as evidenced by the high efficiency of deep-learning classifier to identify ALS MN cultures uniquely based on βIII-tubulin fluorescent images. We also show modest (albeit significant) perturbations in the nuclear compartment given the predictive value of the DAPI fluorescent images in identifying ALS MN cultures. While it remains unclear whether these are strictly pathogenic events, the similar phenotypes detected in the neuritome of MN cultures exposed to heat stress suggests that these events relate to a form of MN stress. Through a thorough comparison of heat, oxidative and osmotic stress induced changes in both cellular shape and ALS-related RBP immunolabeling, we further demonstrate that neuritome-associated perturbations were also detected in control MNs cultured in three different stress conditions. These findings support the notion that the neuronal processes exhibit large perturbations across various stress conditions and argue for increased focus on this cellular subcompartment in future research. Another striking finding is the correlation between recovery kinetics of the neuritome compartment after osmotic and heat stresses, and those of several RBP-related phenotypes. Assuming that the RBP-related phenotypes captured by the CNN-based classifier relate to an RBP change in cellular localization, this result suggests that previously observed stress-induced RBPs mislocalizations are coupled to global changes in the neuritome ^16^. Indeed neurite degeneration has been shown to occur upon oxidative stress through the cytoplasmic sequestration of two proteins (PRMT1 and Nd1-L) in in vitro models of FUS mutant-related ALS ^35^. Furthermore TDP-43 mislocalization and aggregation has also been demonstrated in dystrophic neurites ^1, 28^, while we recently reported an increase in wild-type FUS within neuronal processes in VCP-mutant motor neurons ^30^. Finally, a regulatory role for FUS has also been shown in synaptic formation and function ^35–38^ and aberrant FUS activity in the axonal compartment has been evidenced in a FUS mutant ALS mouse model ^39^. Altogether these studies support the hypothesis of an association between RBP mislocalization and aberrant neuronal processes in ALS. The finding that several RBP-related phenotypes present similar recovery patterns as the neuritome further suggests that additional ALS-related RBPs might exhibit similar aberrant neurite localization in ALS. Future work will directly address the nature of these perturbations using classic approaches which necessitate nuclear and neurite segmentation and the acquisition of hundreds of measurements from each cellular compartment.

Several lines of evidence support the hypothesis that cellular stress is one central mechanism by which MN death occurs in ALS and *in vitro* models of cellular stress are therefore important tools to investigate ALS disease ^17^. It remains however unknown which type of extrinsic cellular stress most closely approximates ALS pathogenesis, and relatively little is known about the effect of thermal stimulation, hyperosmolarity or oxidative stress on the neuritome compartment. Here we find that iPSC-derived MNs exposed to heat stress, as opposed to hyperosmolarity or oxidative stress, closely recapitulate the phenotypes of ALS MN cultures captured by several classifiers. This result suggests that heat stress more closely approximates ALS pathogenesis compared to osmotic and oxidative stressors, which is in line with previous studies in yeast implicating heat stress as being relevant to the study of neurodegeneration ^40^. In particular, our study demonstrates the ability of heat stress to induce subtle ALS-related cellular changes associated within the neuritome compartment and within the FUS fluorescent images. Interestingly we previously found that heat stress alone caused cell death in a iPSC-derived model of MNs ^16^. Furthermore, heat stress ^41, 42^, ageing and neurodegeneration ^43–45^ all associate with intron retention ^2, 4, 46, 47^. Future research will investigate the molecular mechanisms by which elevated temperatures lead to similar responses as those observed in patient-specific ALS MN cultures.

## MATERIALS AND METHODS

### Compliance with ethical standards

Informed consent was obtained from all patients and control controls in this study. Experimental protocols were all carried out according to approved regulations and guidelines by UCLH’s National Hospital for Neurology and Neurosurgery and UCL’s Institute of Neurology joint research ethics committee (09/0272). Cell culture, stress treatments, immunohistochemistry and image acquisition were performed as in ^16^. Indeed these data are utilised in the current manuscript and no additional experiment was required.

### High-content Imaging Dataset

The imaging dataset used in this study consists of fluorescence microscopy images of iPSC-derived motor neurons as previously reported ^16^. The neurons either came from control cell lines or cell lines with the ALS-related VCP mutation and underwent experimentation after six days of terminal differentiation. Details of iPSC lines are provided in **Table S1**. To induce stress the cultures were subject to one hour of oxidative stress, one hour of osmotic stress and one hour of heat stress. To examine recovery, the cultures were subject to one hour of stress and then returned to untreated conditions for two hours following heat stress and one, two and six hours following osmotic stress. Following stress treatments or recovery, cultures were fixed and then immunostained with a combination of three markers, specifically a nuclear-specific marker (DAPI), a neuron-specific marker allowing to outline the neurites (βIII-tubulin) and an antibody against TDP-43, SPFQ, FUS, hnRNPA1 or hnRNPK. The dataset is divided in different experiments (repeats done different days), each with several 96-well plates. Each well corresponds to one cell line, one stress condition and one combination of fluorescent markers. Each well has several non-overlapping fields of view (ranging from 10 to 12) and each field of view has several planes or z-stacks (ranging from 3 to 5) with 1µm steps, generating a large-scale imaging dataset of 156,577 images (**Figs. 1A,B**). The data-set will be deposited on IDR in a close future.

### Image Pre-Processing

All images went through preprocessing steps described in **Supplementary Figure 1.** Raw images are 16-bit images. 16-bit raw z-stack images (1080 x 1080 pixels) from the same field of view were first merged using Maximum Intensity Projection, where the pixel with maximum intensity across all z-stacks is selected at each location in the image. Following conversion of MIP images to 8-bit images, channels were merged together to form an RGB image. We created 13 types of RGB images, either composed of one, two or three channels, to train image classifiers with 13 different combinations of immunostained images (**Figure 1E**). For images with three channels, DAPI was assigned to blue channel, βIII-tubulin to the red channel and the RBP to green channel. For images with one or two channels, pitch-black images were assigned to the remaining channels so that the image would still be considered RGB. Images were then enhanced using Python Image Library Pillow ImageOps ^48^ auto contrast function, to normalize image contrast. This function calculates a histogram of the input image, removes 0.1% of the lightest and darkest pixels from the histogram, and remaps the image so that the darkest pixel becomes black (0), and the lightest becomes white (255). In the fourth step, the enhanced images were divided into 16 smaller images of size 270 x 270 pixels, which allowed better resolution and more images. Structures at this scale proved to be more distinguishable with integrated gradients, and yielded similar results than with whole images. This division also made sense for the fifth step, which consisted in resizing images to 224 x 224 pixels. Finally, images were normalized using mean intensity = [0.485, 0.456, 0.406] and std = [0.229, 0.224, 0.225] across the images from the ImageNet dataset. The last two steps were added in order to fulfill the requirements when using pre-trained models, which expect input images to be normalized in the same way as the dataset on which they were trained.

### Data augmentation

In order to improve accuracy and reduce overfitting, we performed five augmentations on each image of the training set as follows and as previously described ^49^: 1) 90 degrees rotation, 2) one horizontal mirror, 3) one vertical mirror, 4) 90 degrees rotation of horizontal mirror and 5) 90 degrees rotation of vertical mirror. This results in a 6-fold increase of the number of available images for training (five rotations + original).

### CNN-based Image Classifiers Training

We trained 52 Convolutional Neural Network (CNN)-based classifiers to discriminate 1) ALS from control MNs cultures (isALS test), untreated MNs cultures from MNs cultures exposed to 2) oxidative (isOXIDATIVE test), 3) heat (isHEAT test) and 4) osmotic stress (isOSMOTIC test) using 13 combinations of RGB images composed of different channels: either a single channel was used (DAPI, BIII or RBP), either two channels (DAPI:BIII) or three channels (DAPI:BIII:RBP), pitch-black images being assigned to the unused channels (**Table S2**). Instead of training a full new neural network, we performed transfer learning which can be used to address the relatively small number of available images by introducing information from another domain ^50^. For training, images were fed into torchvision MobileNetV2 model which has been pre-trained on ImageNet ^51, 52^. MobileNetV2 is a CNN based on a streamlined architecture that uses depth-wise separable convolutions to build lightweight deep neural networks and which is effective for fine-grained image classification. MobileNetV2 is a lightweight neural network of 3.5 millions parameters, as opposed to the widely used ResNet which contained 11.7 millions parameters, making it suitable for fine-tuning with limited number of images ^53^. All layers of the pre-trained CNN classifier were fine-tuned on our dataset, allowing the training of a highly accurate model with a relatively small training dataset ^54^. The last layer was modified so that it turned the features into predictions for two classes instead of the thousand classes from ImageNet. Training was performed by stochastic gradient descent with learning rate 0.001, batch size 32, using the cross entropy loss function. The training was stopped after 10 epochs. A 10-fold cross validation scheme has been used to evaluate the accuracy of the classification predictions generated by the trained classifiers: the images were shuffled randomly and divided into 10 stratified folds, preserving the percentage of samples for each class; each fold was used once as a test dataset, while the remaining folds were used for the training dataset. Receiver operating characteristic (ROC) curves were generated to evaluate the model’s ability to distinguish two cell culture conditions. Receiver operating characteristics (ROC) curves plot the true positive rate (sensitivity) versus the false positive rate (1 – specificity). The area under the ROC curve was used as the performance measure or classification accuracy. The classification accuracy over all folds is reported in **Table S3**. We evaluated the 52 trained models on all MN cultures when the right combination of markers were available (typically FUS, DAPI and BIII are available for some MNs cultures while SFPQ, DAPI and BIII are available for others). The probabilities to belong to one of the four tested conditions (iALS, isOXIDATIVE, isOSMOTIC, isHEAT) outputted by the classifiers are then aggregated by computing the average probability over all of the 16 cropped images originating from a single image, thereby obtaining a single probability per original-sized images. A single probability per MN culture is reported for each of the fours test by averaging the signal over all images (typically 7 per MN culture) as reported in **Tables S4-S7**.

### Model Explainability and Integrated Gradient

The integrated gradient (IG) is a widely used interpretability algorithm that allows to identify what pixels of an image have the strongest effect on the model’s predicted class probabilities and therefore allowing to visualize which parts or the image are important for classification ^26^, by computing the gradient of the model’s prediction output to its input features. We used the Captum Insights method ^55^ to obtain the IG for randomly selected images associated with high classifier prediction scores.

### Model prediction data analysis

We used R and lme4 ^56^ to perform linear mixed effects analysis of the relationship between individual model predictions and either stress MNs culture condition (at the corresponding time after treatment) or VCP-mutant MNs cultures, accounting for idiosyncratic variation due to either cell line or experiments. As fixed effects, we either entered the stress condition or the VCP-mutation variable into the model. As random effects, we had intercepts for either cell lines (CTRL1, CTRL2, CTRL3, MUT1, MUT2, MUT3, MUT4) and experiments. P-values were obtained by likelihood ratio tests of the full model with the effect in question against the model without the effect in question, i.e. comparing a full linear model fitting the classifier predictions using both the fixed effect (the stress or disease) and the random effects (cell line and experiments), with a reduced linear model, which only considers the random effects.

### Data and software availability

We provide raw images, complete source code and trained models to readily reproduce figures, tables, and other results that involve computation in order to facilitate the development and evaluation of additional profiling methods. The image data that support the findings of this study will be uploaded to the Image Data Resource ^57, 58^. As this requires some more time given the size of the data (1TB), we have uploaded a subset of the data on Zenodo under the accession number 4664177. The 52 trained deep learning models have been uploaded on Zenodo under the accession number 4664252. The code for obtaining the models as well as the Jupyter notebooks are available at https://github.com/idiap/als-classification.

## AUTHOR CONTRIBUTIONS

Conceptualization, R.L., R.P.; Formal Analysis, C.V., R.L.; Investigation, C.V., J. H.; Writing – Original Draft, R.L., R.P.; Writing – Review & Editing, R.L., R.P., C.V., J.H.; Resources, R.P.; Visualization, C.V., J.H..; Funding Acquisition, R.L., R.P.; Supervision, R.L., R.P.

## ACKNOWLEDGMENTS

This work was supported by the Idiap Research Institute and by the Francis Crick Institute which receives its core funding from Cancer Research UK (FC010110), the UK Medical Research Council (FC010110), and the Wellcome Trust (FC010110). R.P. holds an MRC Senior Clinical Fellowship [MR/S006591/1]. We thank Olivier Bornet, Bastien Crettol and Philip Abbet for their technical support in generating the repositories for the code, the models and the dataset.

## Electronic supplementary material

Supplementary Tables 1-7 can be accessed <u>here</u>.

Table S1 | Description of human sample origin and mutations.

Table S2 | List of the ALS and stress trained CNN-based classifiers.

Table S3 | Performances of the 52 trained classifiers across the 10 folds.

Table S4 | ALS classifier assigned class probabilities for all the views from a cell culture (∼ 10 per cell culture) that are then averaged to obtain a single per-culture classification probability.

Table S5 | Same as Table S4 for oxidative stress classifier.

Table S6 | Same as Table S4 for heat stress classifier.

Table S7 | Same as Table S4 for osmotic stress classifier.

---

**Supplementary Figure 1.**
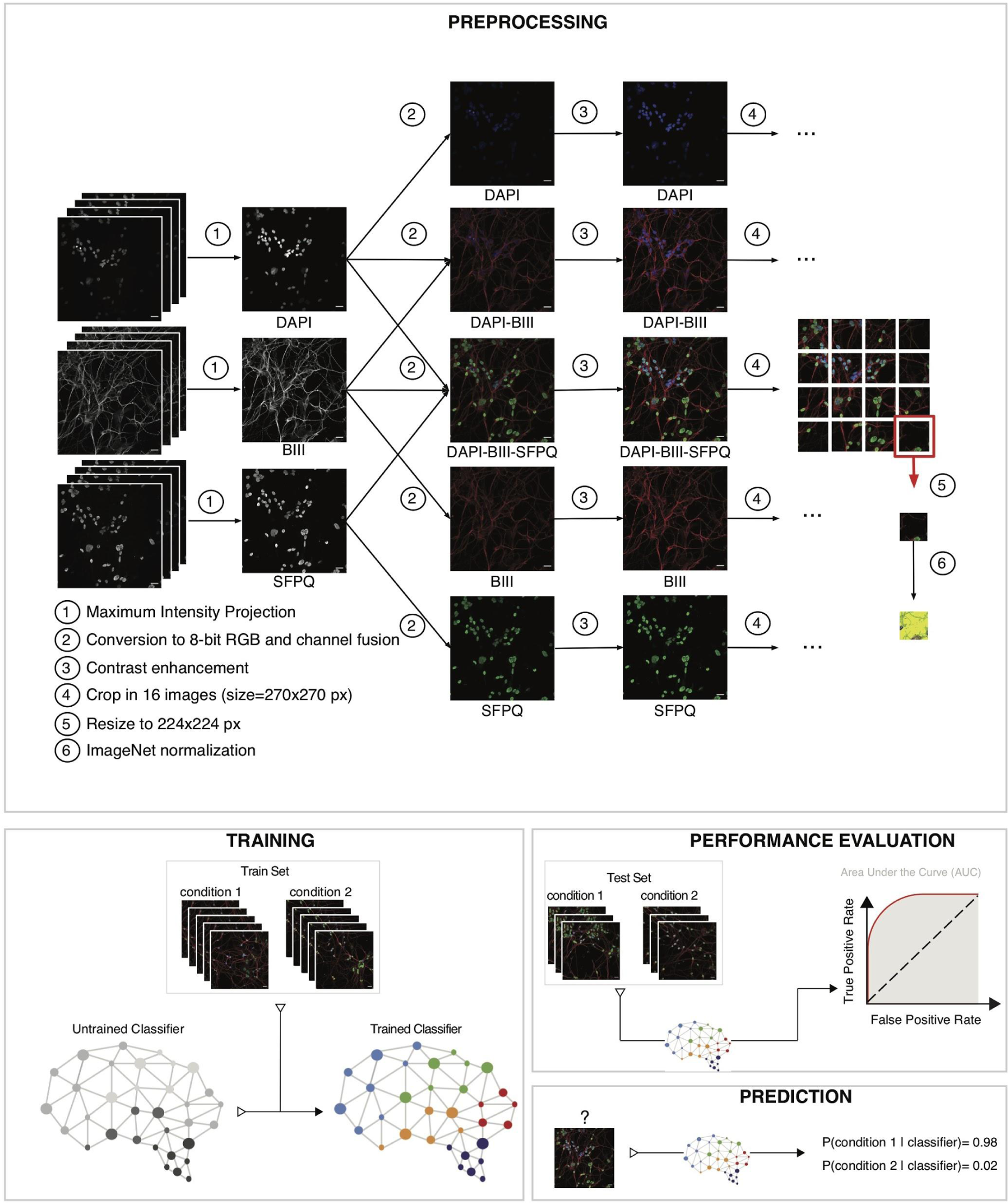
Schematic depicting the CNN-based approach. **Preprocessing:** (1) 16-bit z-stack raw images are merged using the Maximum Intensity Projection (MIP); (2) the MIP images are then converted to 8-bit images and the different channels are merged together, depending on the protocol (either 1, 2 or 3 channels); (3) contrast is enhanced with a cutoff of 0.1%; (4) each image (size=1080x1080 pixels) is divided into 16 images of 270x270 pixels; (5) images are then resized into 224x224 pixels; (6) images are normalized the same way as the ImageNet dataset, using mean = [0.485, 0.456, 0.406] and std = [0.229, 0.224, 0.225]. **Training:** Concept illustration of training a neural network for binary classification of one condition against another. The train set is composed of labeled images for both conditions and is used to train the network, which assigns more weight to discriminating features in the images. **Performance Evaluation:** A distinct set of labeled images, the test set, is used to evaluate the performance of the trained network using metrics such as the Area Under the receiver operating characteristic Curve (AUC). The performances of different classifiers trained with different markers can be compared to uncover the relevance of specific markers in discriminating two conditions. **Prediction:** The trained model can then evaluate the probability for a given unlabeled image to belong to either one or the other condition.

**Supplementary Figure 2.**
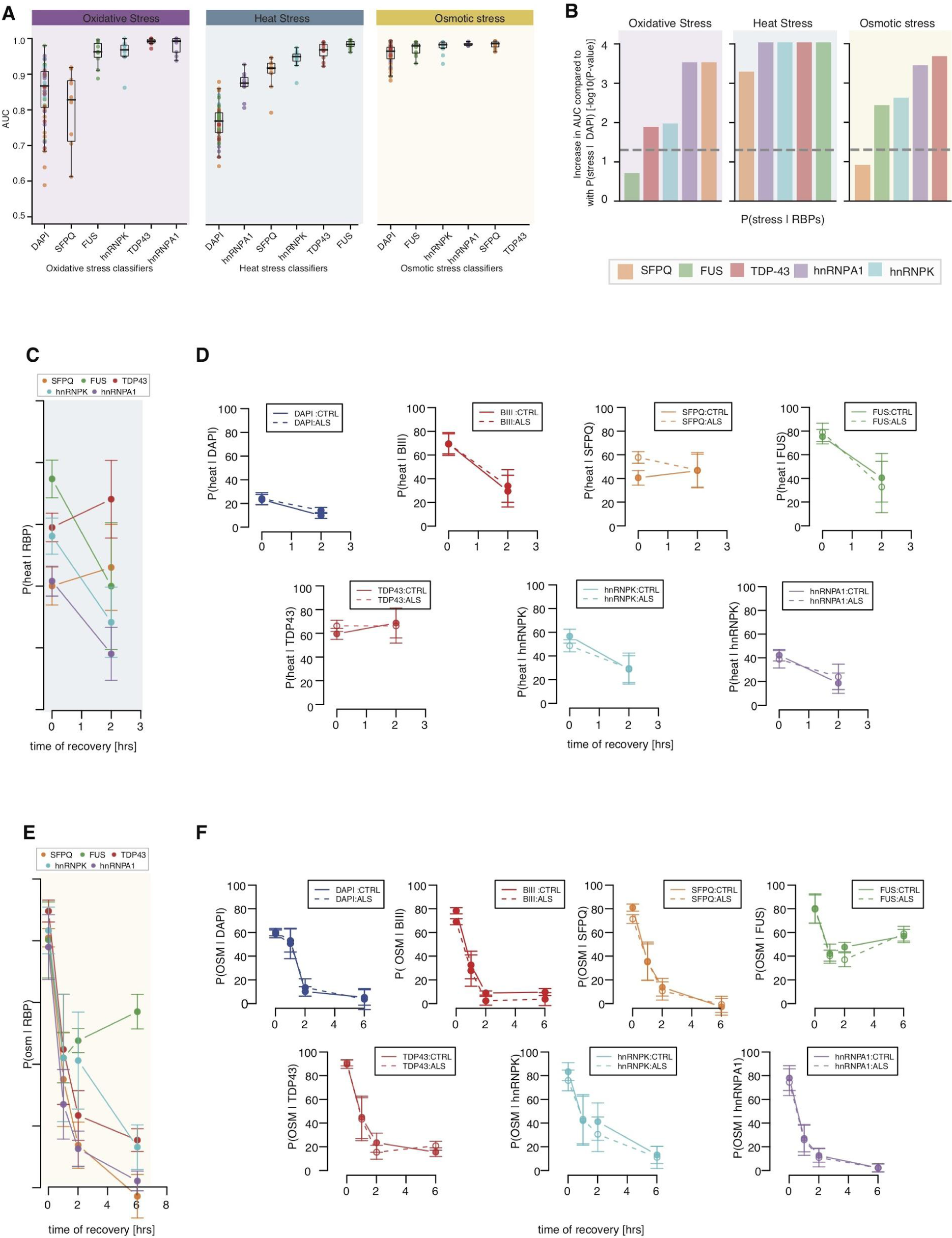
(**A**) Boxplots showing the distributions of model performances as evaluated using the AUC for classifiers trained using DAPI or ALS-related RBP markers to discriminate untreated from stressed MN cultures. Each classifier was submitted to 10-fold cross-validation in 5 different subsets of the data, resulting in 50 points per classifier. Boxplots display the five number summary of median, lower and upper quartiles, minimum and maximum values. P-values obtained from a one-sided Mann-Whitney test. (**B**) Bar graphs representing the increase in performance as obtained from -log10(P-values) of one-sided Mann-Whitney test comparing the AUCs from the *stress|DAPI* classifier and the AUCs from individual *stress|DAPI:RBP* classifiers for oxidative, heat and osmotic stresses. (**C**) Effect size (mean ± standard errors) of *heat|RBPs* classifier predictions of control MN cultures one hour after heat stress and two hours after recovery from heat stress. Effect size of the treatment at each time-point is obtained using linear mixed effects analysis accounting for idiosyncratic variations due to cell lines and experiment bias. (**D**) Same as (C) for individual *heat|RBPs* classifiers predictions. Solid lines = control MN cultures. Dashed lines = VCP-mutant MN cultures. (**E**) Effect size (mean ± standard errors) of *osm|RBPs* classifiers predict control MN cultures one hour after osmotic stress and one, two and six hours after recovery from osmotic stress. (**F**) Same as (E) for individual *osm|RBPs* classifiers predictions. Solid lines = control MN cultures. Dashed lines = VCP-mutant MN cultures.

**Supplementary Figure 3.**
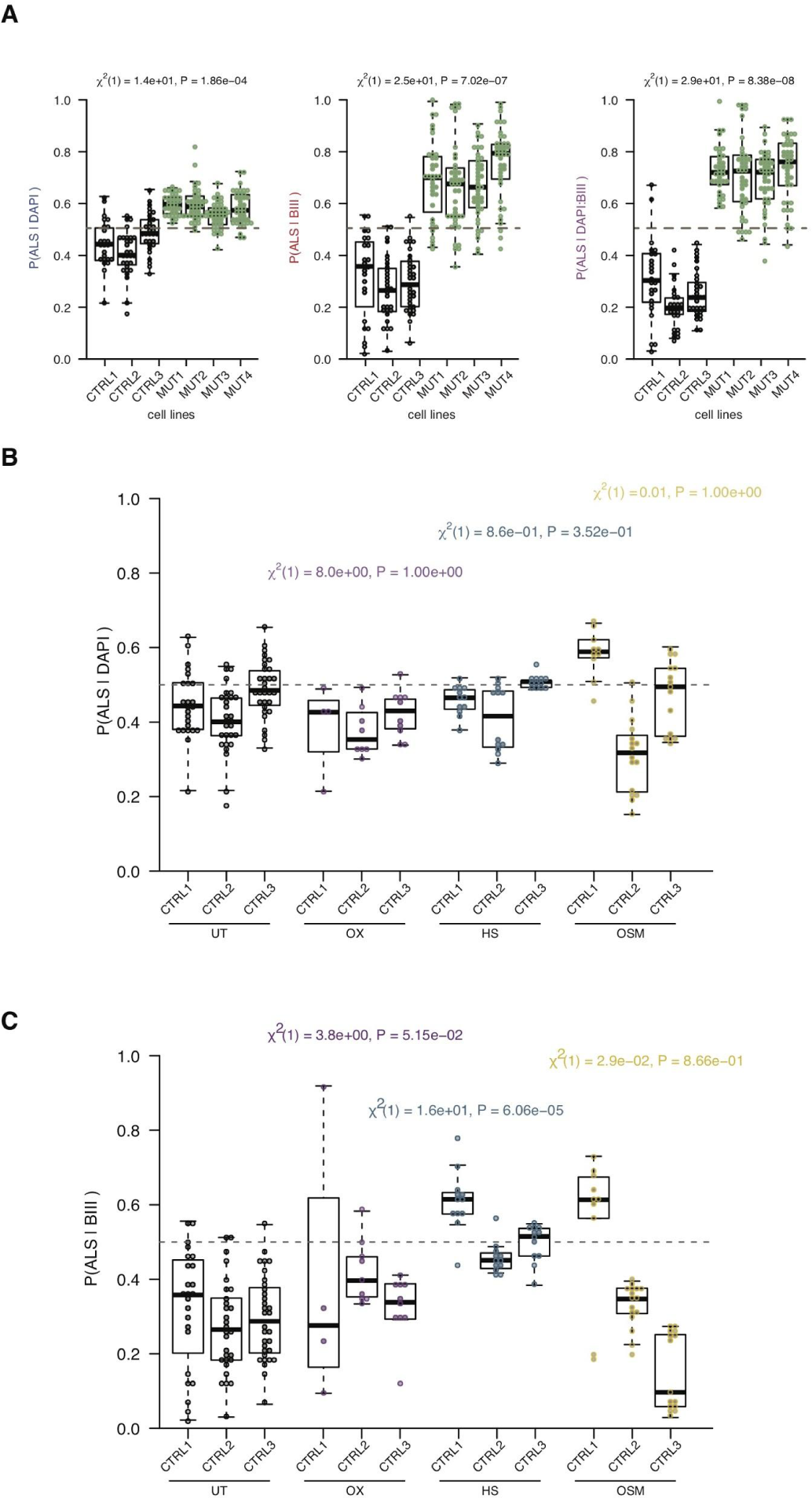
(**A**) Distributions of the *ALS|DAPI* (*left*), *ALS|BIII* (*center*) and *ALS|DAPI:BIII* (*right*) model predictions for the individual MN cultures originating from 3 control cell lines (*grey dots*) and 4 VCP-mutant cell lines (*green dots*). Linear mixed effects analysis of the relationship between each model prediction and VCP mutation to account for idiosyncratic variation due to cell line or experiment differences. VCP mutation significantly increases *ALS|DAPI* predictions [χ^2^(1) = 14 and *P* = 2.28*e*^−04^] by about 0. 13 ± 0. 023 (standard errors), *ALS|BIII* predictions [χ^2^ (1) = 23 and *P* = 2. 1*e*^-06^] by about 0. 4 ± 0. 033 (standard errors), and *ALS|DAPI:BIII* predictions [χ^2^ (1) = 29 and *P* = 8. 4*e*^-08^] by about 0.47 ± 0. 031 (standard errors). (**B**) Distributions of the *ALS|DAPI* model predictions for the individual MN cultures originating from three control cell lines after one hour of oxidative (magenta), heat (blue) and osmotic (yellow) stress. Linear mixed effects analysis of the relationship between each model prediction and individual treatment effect to account for idiosyncratic variation due to cell line or experiment differences. (**C**) Distributions of the *ALS|BIII* model predictions for the individual MNs cultures originating from 3 control cell lines after one hour of oxidative (magenta), heat (blue) and osmotic (yellow) stress. Linear mixed effects analysis of the relationship between each model prediction and individual treatment effect to account for idiosyncratic variation due to cell line or experiment differences.

**Supplementary Figure 4.**
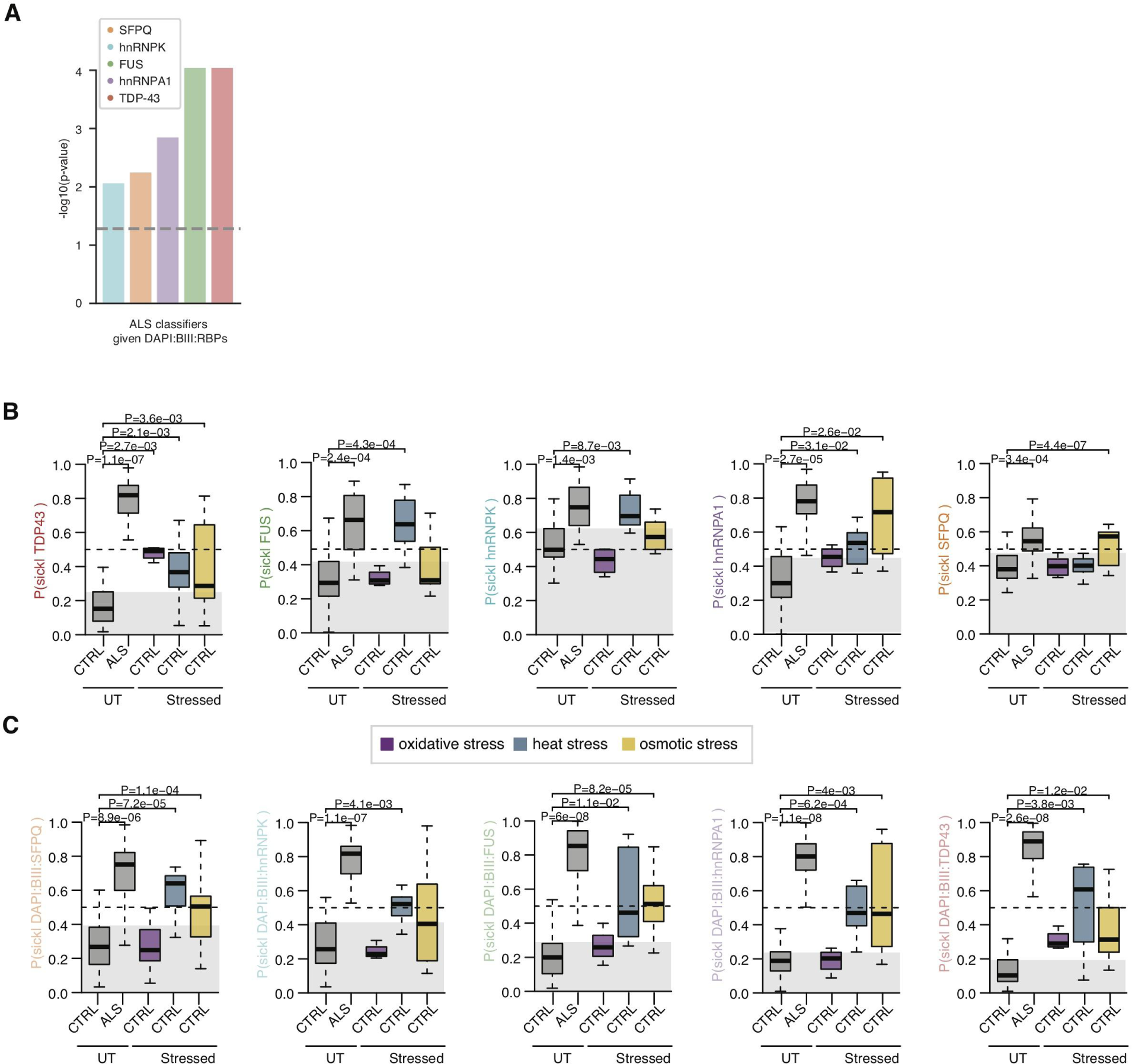
(**A**) Bar graphs representing the increase in performance as obtained from -log10(P-values) of one-sided Mann-Whitney test comparing the AUCs from the *ALS|DAPI:BIII* classifier and the AUCs from individual *ALS|DAPI:BIII:RBP* classifiers. (**B**) Boxplots showing the distributions *ALS|*RBPs model predictions on untreated control and ALS MN cultures, and control MN cultures after one one hour of oxidative, heat and osmotic stress. Magenta = oxidative stress. Blue = heat stress. Yellow = osmotic stress. Stress treatment effect analysis on model prediction obtained using linear mixed effects analysis. P-values obtained from linear mixed models are indicated when significant. (**C**) Same as (B) for *ALS|DAPI:BIII:*RBPs model predictions.

